# BlastPhyMe: A toolkit for rapid generation and analysis of protein-coding sequence datasets

**DOI:** 10.1101/059881

**Authors:** Daniel Gow, Alexis Ballesteros, Belinda SW Chang, Ryan K Schott

## Abstract

**Summary:** We present BlastPhyMe (BLAST, Phylogenies, and Molecular Evolution) a new application to facilitate the fast and easy generation and analysis of protein-coding sequence datasets. The application uses a portable database framework to manage and organize sequences along with a graphical user interface (GUI) that makes the application extremely easy to use. BlastPhyMe utilizes several existing services and applications in a unique way that saves researchers considerable time when building and analyzing protein-coding datasets. The application consists of two modules that can be used separately or together. The first module enables the assembly of coding sequence datasets. BLAST searches can be used to obtain all related sequences of interest from NCBI. Full GenBank records are saved within the database and coding sequences are automatically extracted. A feature of particular note is that sequences can be sorted based on NCBI taxonomic hierarchy before export export in formats such as FASTA for visualization using existing tools, such as MEGA. The application provides GUIs for automatic alignment of sequences with the popular tools MUSCLE and PRANK, as well as for reconstructing phylogenetic trees using PhyML. The second module incorporates selection analyses using codon-based likelihood methods. The alignments and phylogenetic trees generated with the dataset module, or those generated elsewhere, can be used to run the models implemented in the codeml PAML package. A GUI allows easy selection of models and parameters. Importantly, replicate analyses with different parameter starting values can be automatically performed in order to ensure selection of the best-fitting model. Multiple analyses can be run simultaneously based on the number of processor cores available, while additional analyses will be run iteratively until completed. Results are saved within the database and can be exported to publication-ready Excel tables, which further automatically compute the appropriate likelihood ratio test between models in order to determine statistical significance. Future development will expand the available phylogenetic and molecular-evolutionary analyses. BlastPhyMe saves researchers of all bioinformatics experience levels considerable time by automating the numerous tasks required for the generation and analysis of protein-coding sequence datasets using a straightforward graphical interface.

**Availability:** Installation package and source code available from: https://github.com/Schott-Lab/BlastPhyMe

## Introduction

With continued advances in Next Generation Sequencing (NGS) technology and ongoing genome sequencing projects (e.g., Vertebrate Genomes Project, Formenti et al. 2026; Earth BioGenome Project, Blaxter et al. 2025) the amount of publicly available sequence data is growing exponentially. This data provides a valuable resource for biologists, but as the number of available sequences continues to grow the generation and analysis of comparative sequence datasets has become a daunting task, especially for researchers that lack bioinformatics and scripting experience. While databases such as GenBank provide access to the vast number of publicly available sequences, searching for, finding, and extracting sequences from GenBank through a web browser is extremely time consuming. Additionally, many bioinformatics tools for the alignment and analysis of sequences are command-line based, which can be difficult to use, produce outputs that are difficult to understand, and are time consuming to manually parse. To address these difficulties, we have developed **BlastPhyMe** (BLAST, Phylogenies, and Molecular Evolution), a new application that automatically gathers, organizes, and analyzes gene sequences within a portable database framework using an intuitive graphical user interface (GUI).

BlastPhyMe consists of two modules that can be used separately or together: Gene Sequences and Selection Analyses (PAML). The Gene Sequences module enables the assembly of coding sequence datasets. This includes searches and automatic extraction of sequences from GenBank using direct and BLAST searches, import of user data, multiple sequence alignments, and phylogenetic analyses. The Selection Analyses (PAML) module incorporates selection analyses using codon-based likelihood methods. The alignments and phylogenetic trees generated with the Gene Sequences module, or those generated elsewhere, can be used to run the models implemented in the codeml PAML package using a series of GUIs. Results are saved within the database and can be exported to publication-ready Excel tables, which further automatically compute the appropriate statistical tests to evaluate model significance.

## Methods

BlastPhyMe is a Microsoft Windows application coded in the C# programming language on the Microsoft .NET Framework platform, version 4.8.1 (https://learn.microsoft.com/en-us/dotnet/framework/). The database engine that BlastPhyMe interacts with for storing data is the Microsoft SQL Server 2019 Express LocalDB engine (https://learn.microsoft.com/en-us/sql/database-engine/configure-windows/sql-server-express-localdb?view=sql-serverver15). For communicating with the NCBI BLASTN web service, BlastPhyMe uses the .NET Bio open-source library (https://github.com/dotnetbio/bio). BlastPhyMe has been designed to the standard of a 3-Tier architecture. The application code is separated between three distinct layers: user interface, middle tier, and database (Figure 1). The software is organized in a Visual Studio solution file containing three C# projects (.csproj): Pilgrimage, PilgrimageTests, and ChangLab. The C# project PilgrimageTests is used for software development purposes meant to simulate the user interface during testing.

**Figure 1.**
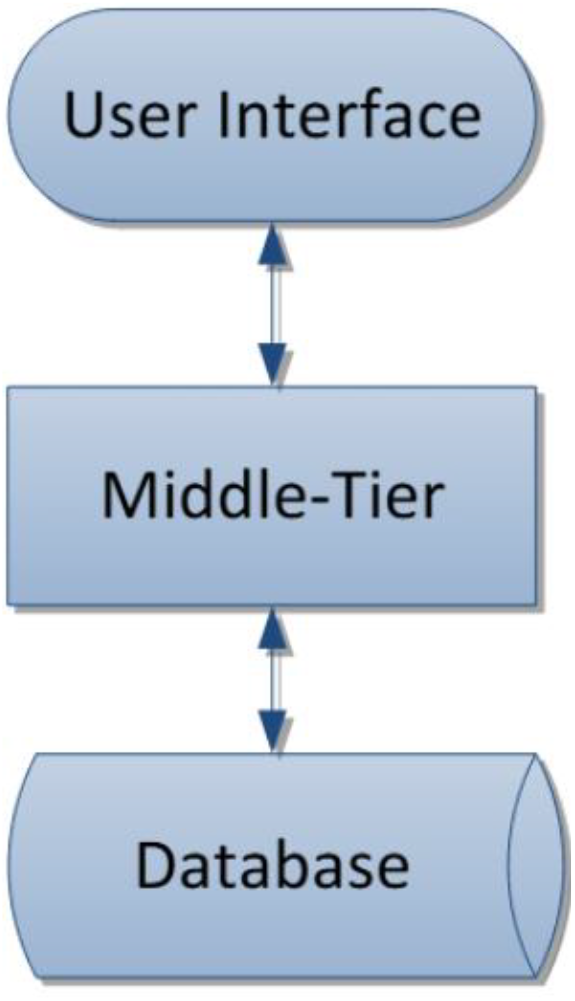
Structure of BlastPhyMe’s three software layers.

### User Interface

The User Interface layer includes all code necessary to display visual interfaces for the user to interact with. User Interface code does not interact directly with the Database layer or third-party systems (e.g., NCBI) and is abstracted from both by objects within the Middle-Tier layer.

### Middle-Tier

The Middle-Tier layer includes all code necessary to transfer data between the User Interface and Database layer. Code within the Middle-Tier interprets the database architecture into objects that are exposed for the User Interface to interact with. The code found in the project is organized into nine folders: BlastN, Common, Genes, Jobs, LocalDatabase, NCBI, PAML, RecordSets, and Taxonomy. The files found in the LocalDatabase folder contain the code needed for the creation and updating of the database layer which consists of .sql and .cs files. The Middle-Tier is responsible for all direct communication with the database, and all interaction with third-party systems and products (e.g., .NET Bio) is contained within the Middle-Tier layer.

### Database Layer

The Database layer comprises all of the architecture necessary to store data for the BlastPhyMe application as well as code to manipulate that data within the database. The BlastPhyMe database exposes stored procedures to handle all data collection and modification processes performed by the Middle-Tier layer. BlastPhyMe is capable of exporting data from its database to a “data file”, distinguished by the “.bpmd” file extension. A BlastPhyMe data file is an XML document compressed via the GZip compression algorithm.

Database migrations are handled by the Autoupdate.cs script, which runs on application startup. The code queries the Common.ApplicationProperty table and compares the database version with the expected version. If a discrepancy is detected, a notification asks the user to update the database file. When the user accepts the prompt, Autoupdate.cs executes the appropriate SQL script to bring the schema up to date.

This ensures that the database layer remains consistent with the application version without manual intervention. The update process is built on Microsoft SQL Server 2019 which is an application used to manage, configure, administer, and develop SQL infrastructure.

### Third-Party Tools

In addition to communicating with NCBI, BlastPhyMe utilizes several third-party programs. These programs are accessed and run through the BlastPhyMe interface. Gene sequences and alignments can be exported to MEGA (Tamura et al. 2013; http://www.megasoftware.net/) for visualization and editing. Multiple sequence alignments can be performed using PRANK (Löytynoja and Goldman 2008; http://wasabiapp.org/software/prank/) and MUSCLE (Edgar 2004; http://www.drive5.com/muscle/). Phylogenetic trees can be inferred using PhyML (Guidon et al. 2010; http://www.atgc-montpellier.fr/phyml/binaries.php). Resulting phylogenetic trees can be sent to Mesquite (https://www.mesquiteproject.org/) for visualization. Selection analyses can be performed using PAML (Yang 2007; http://abacus.gene.ucl.ac.uk/software/paml.html). Finally, sequences and PAML results tables can be exported to Microsoft Excel (www.microsoftstore.com).

BlastPhyMe communicates with NCBI’s GenBank nucleotide database to search for and download GenBank records. This communication is performed via HTTPS requests of the E-utilities web services hosted by NCBI (http://www.ncbi.nlm.nih.gov/books/NBK25499/). BlastPhyMe also communicates with NCBI’s BLAST web service using the aforementioned .NET Bio libraries, which make use of the QBlast URL API (http://www.ncbi.nlm.nih.gov/blast/Doc/urlapi.html).

## Installation

BlastPhyMe comes with a complete installation package available at: https://github.com/ryankschott/BlastPhyMe/releases. Prerequisites will be installed automatically. Third party programs need to be downloaded and installed separately. The software requires Windows 11 or 10 as well as Microsoft .NET Framework 4.8.1 and SQL Server 2019 Express LocalDB engine, which is automatically installed if not present.

Installation proceeds in three steps: (1) download and run BlastPhyMe installer, which automatically installs all prerequisites; (2) download and install third-party programs listed in Table 1; and (3) specify required file paths for third-party executable within the BlastPhyMe software. A complete step-by-step installation guide with screenshots is available in the GitHub repository.

**Table 1.** Required third-party programs.

| **Program** | **Purpose** | **URL** |
|---|---|---|
| **MUSCLE** | Multiple sequence alignments | https://www.drive5.com/muscle |
| **PRANK** | Multiple sequence alignments | https://wasabiapp.org/software/prank/ |
| **PhyML** | Inferring phylogenetic trees | https://www.atgc-montpellier.fr/phyml/binaries.php |
| **PAML (codeml)** | Utilize Selection Analysis module | https://abacus.gene.ucl.ac.uk/software/paml.html. |
| **MEGA** | Visualizing and editing sequences and alignments | https://www.megasoftware.net/ |
| **Mesquite** | Visualize tree files, label foreground branches/clades for the PAML branch, branch-site, and clade models | https://www.mesquiteproject.org/ |
| **Excel** | BlastPhyMe can write results from PAML analyses into Microsoft Excel | https://www.microsoftstore.com/ |

BlastPhyMe’s BLAST search requires an internet connection to communicate with NCBI’s web services. All other features can be run locally without internet access.

## Features and Use

Upon running BlastPhyMe for the first time the user will be asked to create a database. This database will store all of the sequences, analyses, and results that you generate, or import into, BlastPhyMe. The database is stored in a .mdf file. Each user will only need a single database and these can be shared between users. After creating a database file the user will be prompted to create a project. Projects are a way to organize sets of similar data and each database can have multiple projects.

BlastPhyMe consists of two distinct modules: **(1)** Gene Sequences and **(2)** Selection Analyses (PAML). These can be accessed via the tabs as shown in Figure 2. Each module can be used completely independently of the other, but they are also designed to offer a continuous workflow from dataset generation to selection analyses as shown in Figure 3.

**Figure 2.**
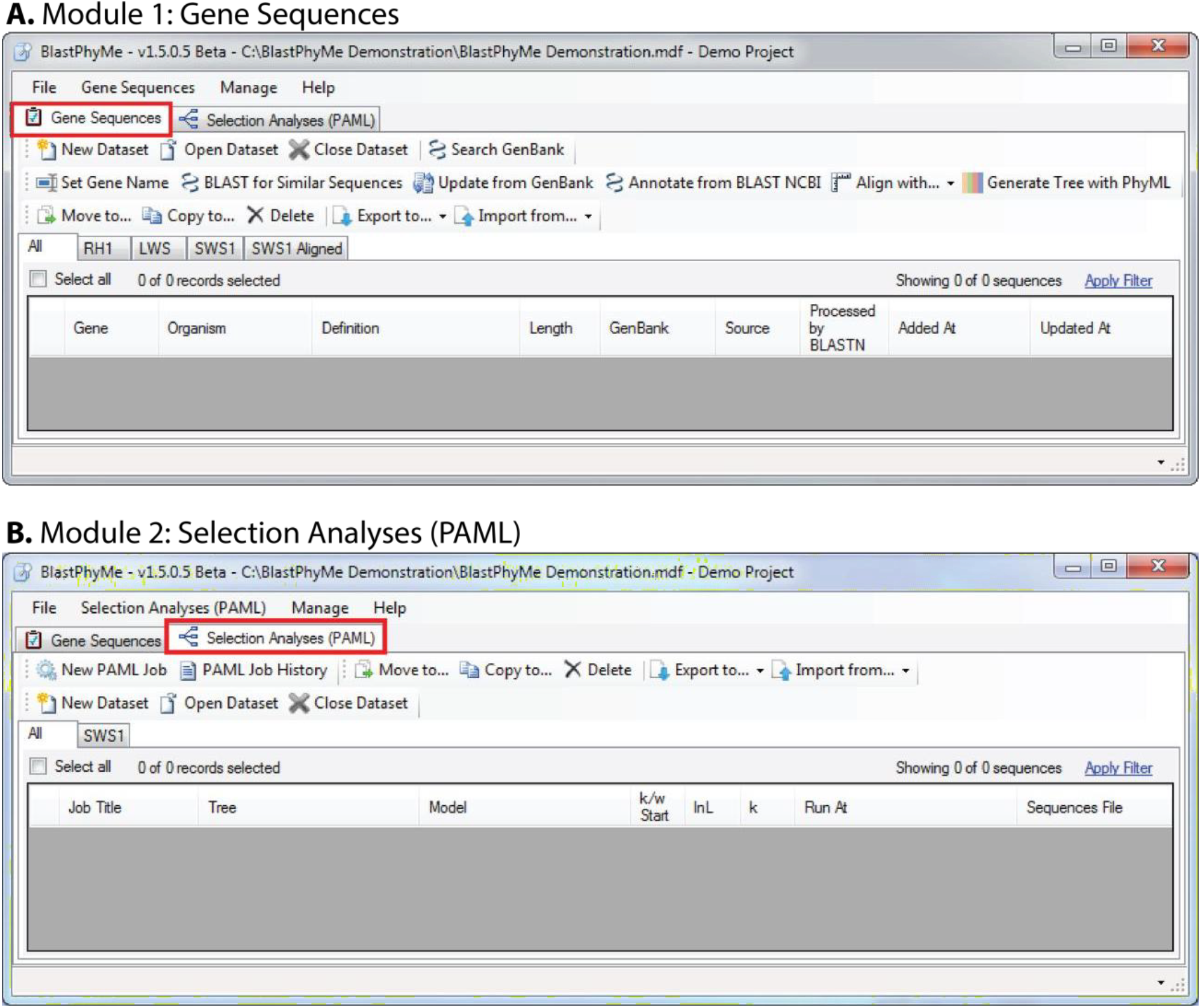
The two distinct modules of BlastPhyMe: Gene Sequences and Selection Analyses (PAML).

**Figure 3.**
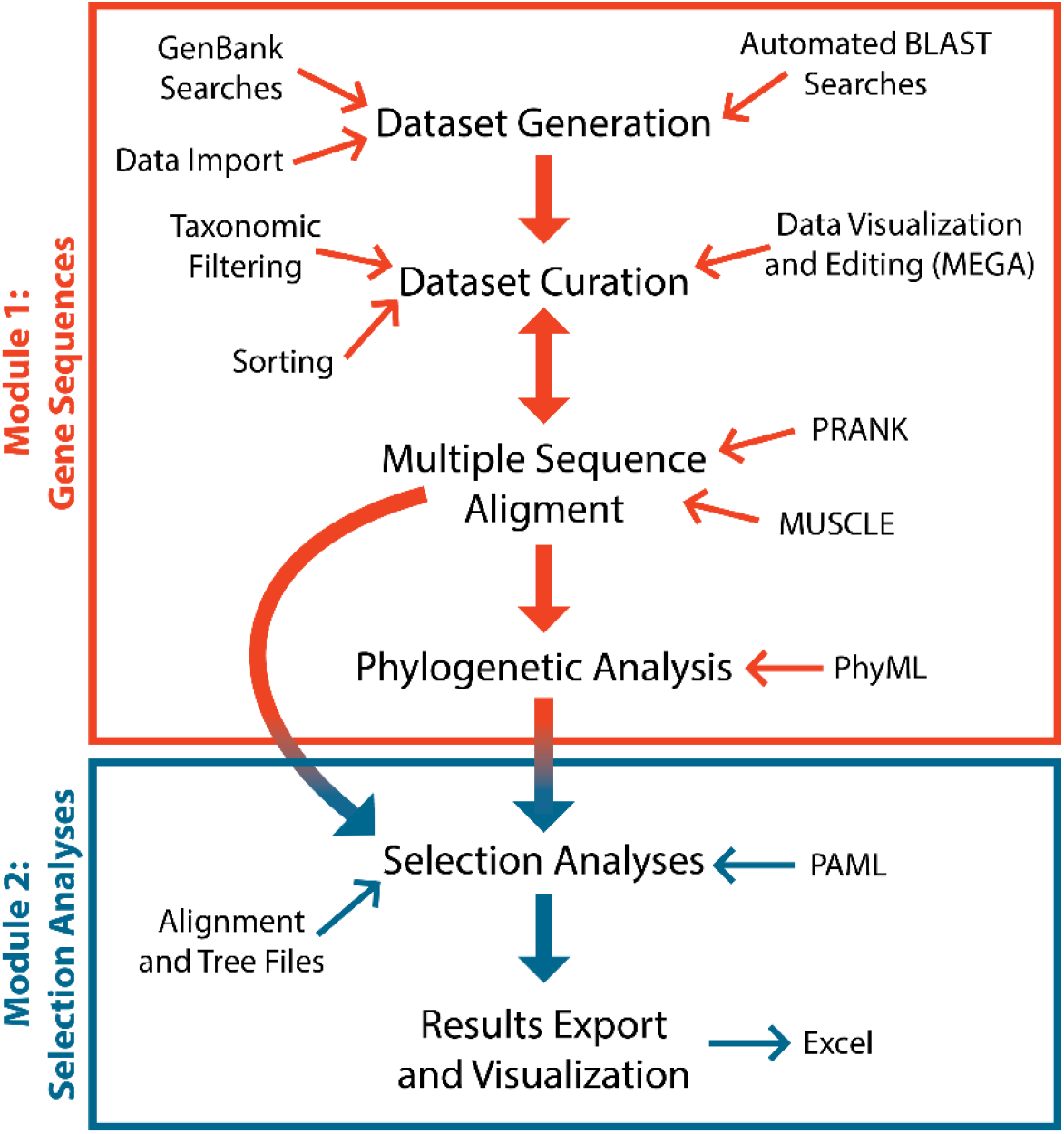
BlastPhyMe workflow. The two modules are distinct, but results from Module 1 provide the necessary input files for Module 2.

### Gene Sequences

The Gene Sequences module is organized into datasets. Each dataset can contain any number of sequences from one or more genes and species. To create a dataset click the new dataset button. Sequences can be added to the dataset using the ‘Search GenBank’ function. User data (e.g., FASTA file) can also be imported using the ‘Import from’ function. The ‘Search GenBank’ function opens a dialog box allowing search terms to be submitted to NCBI GenBank (Fig. 4). All search terms supported by the GenBank website are also supported when submitting with BlastPhyMe. Search results will appear in a separate window (Fig. 5). Sequences that the user wishes to download can be selected and then added to an existing or new dataset using the ‘Add to’ button. This will save the complete GenBank record for the selected sequences.

**Figure 4.**
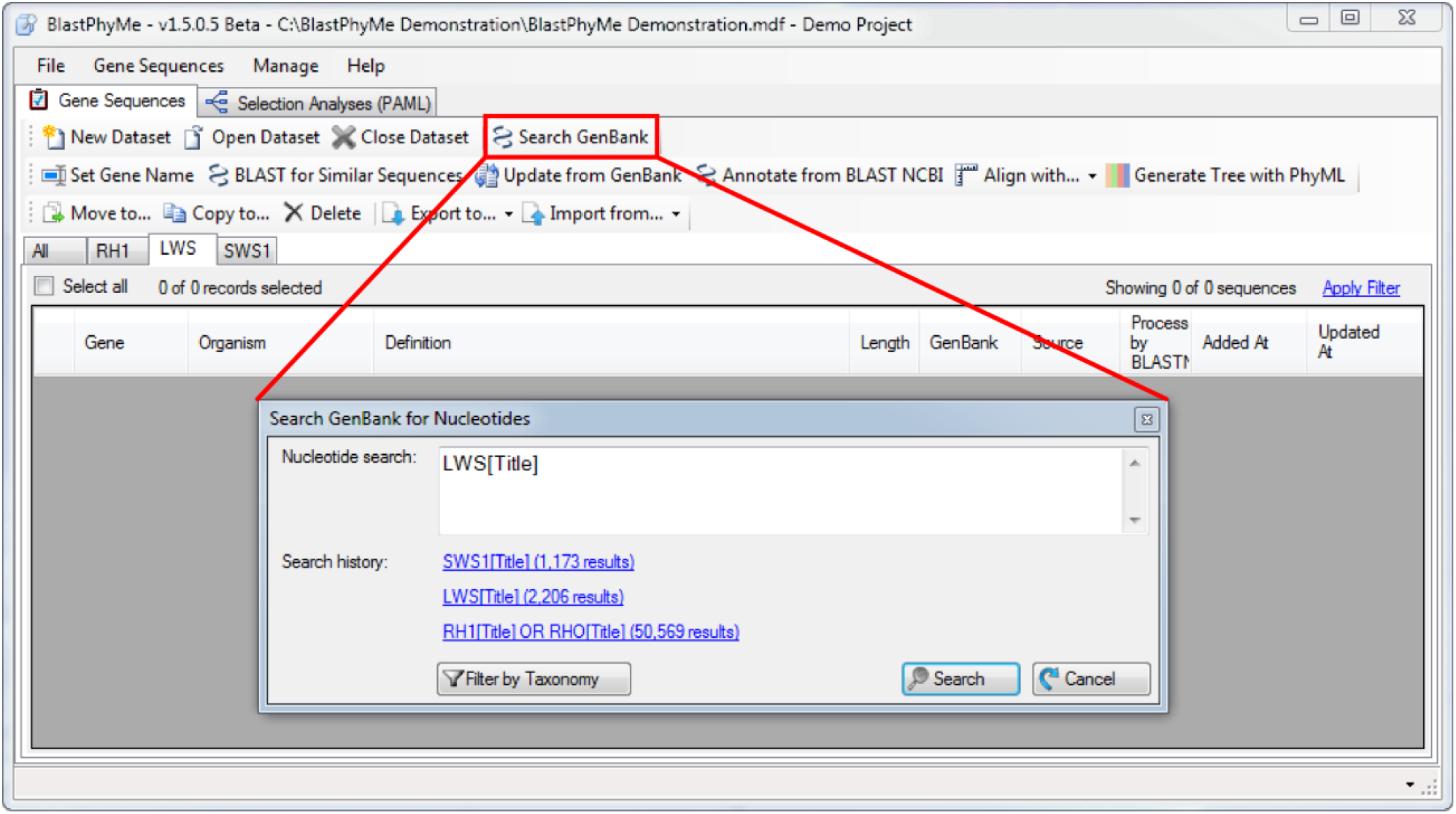
GenBank Search Function. This function directly searches the GenBank nucleotide database. All search terms used on the webpage can be used with BlastPhyMe.

**Figure 5.**
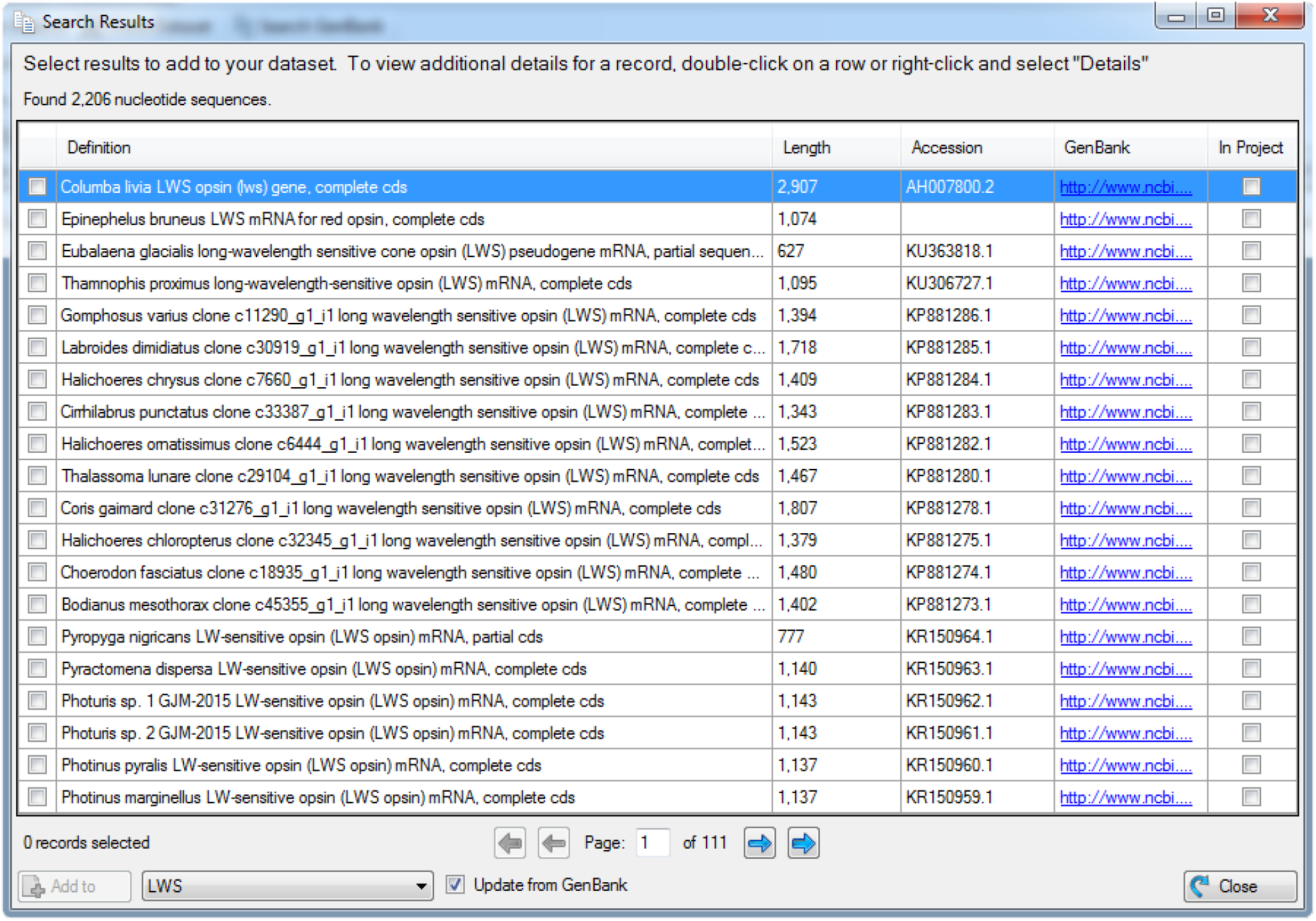
GenBank Search results window. Sequences can be selected using the check boxes and added to a new or existing dataset using the ‘Add to’ button.

Once added to a dataset, sequences can be doubled-clicked to open up a separate window with additional information about the sequence including a link to the GenBank page (Fig. 6). Sequences can be ordered by clicking on any of the column headings. Sequences can be filtered using both text matching and taxonomic filtering using the ‘Apply Filter’ function. Sequences can be deleted and moved or copied to another dataset using the ‘Move to’, ‘Copy to’, and ‘Delete’ buttons. Sequences can be exported to different formats, including FASTA, or opened directly into MEGA for visualization and editing, using the ‘Export to’ button.

**Figure 6.**
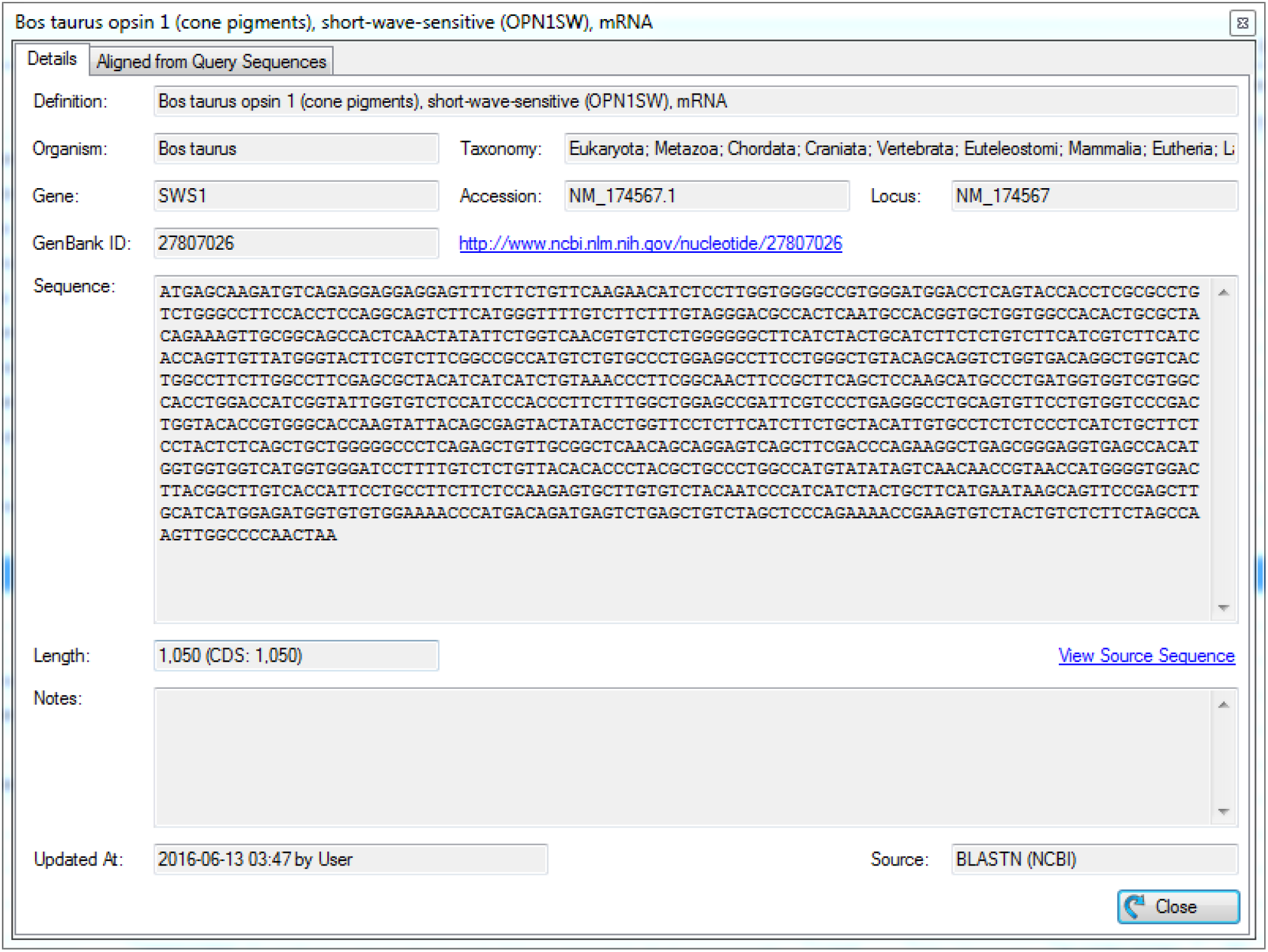
Additional data available for each sequence stored within the database accessible by double clicking on a sequence entry.

The dataset can be further expanded using the ‘BLAST for Similar Sequences’ function. This function will submit selected sequences for BLAST analysis against the full NCBI nucleotide database (Fig. 7). Resulting BLAST hits will be automatically combined, removing duplicates, and can be downloaded and added to a dataset (Fig. 7). This function allows the user to quickly expand a dataset to include all available sequences of a particular gene or set of genes.

**Figure 7.**
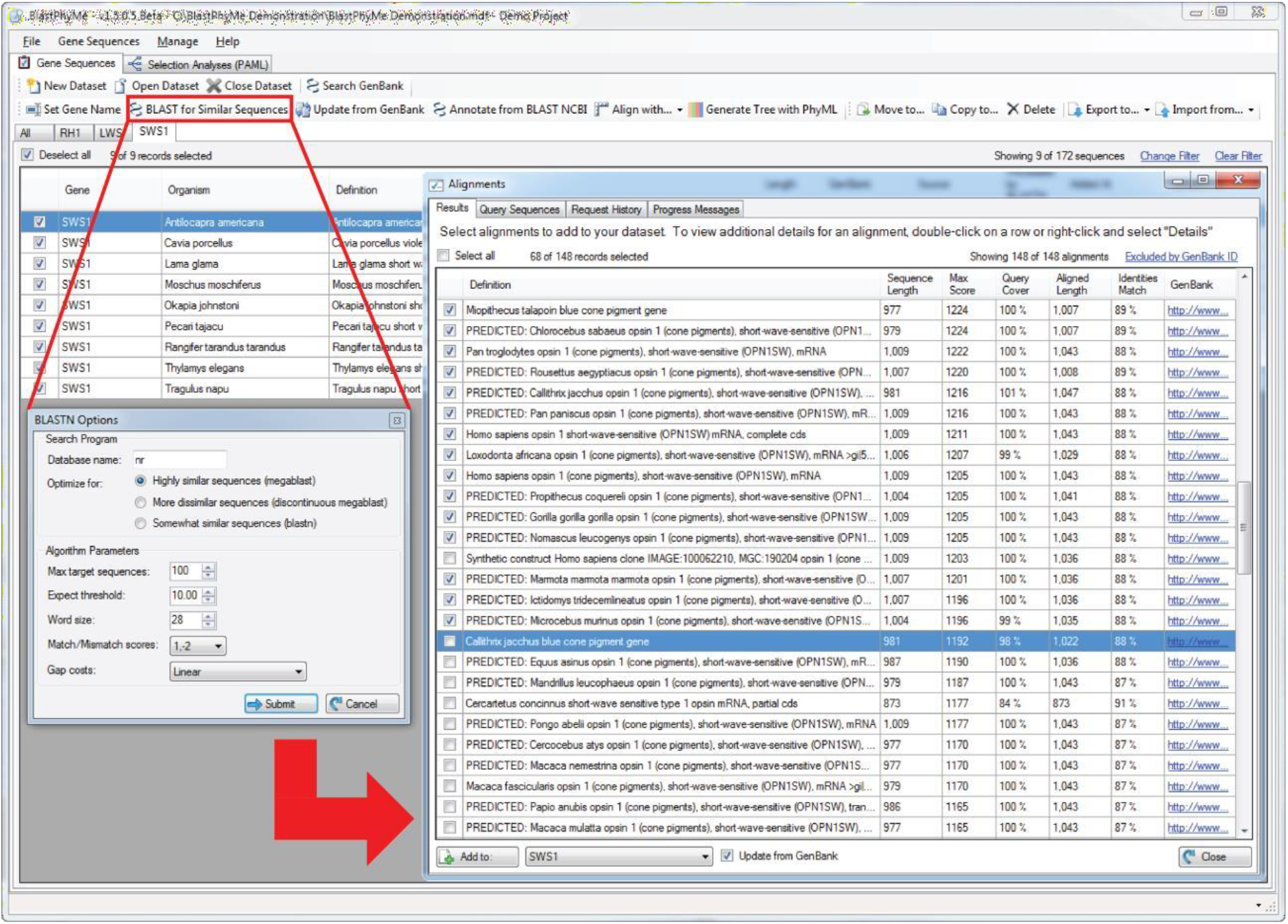
BLASTN search and results windows used to expand a dataset to include additional related sequences from the NCBI nucleotide database.

Selected sequences from a dataset can be aligned using either PRANK or MUSCLE using the ‘Align with’ function. This will open a dialog box allowing parameters to be set (Fig. 8). It will be necessary to add the location of PRANK.exe and MUSCLE.exe on first use. Sequences submitted for PRANK codon alignment will be automatically trimmed to the last complete codon. Upon completion of the alignment a new window will open, allowing the aligned sequences to be selected and added to a new (aligned) dataset. The output of the alignment will also be saved to the selected working directory. The aligned dataset (or an alignment imported separately using the ‘Import from’ function) can then be submitted for phylogenetic analysis using the ‘Generate Tree with PhyML’ function. This function will open a dialog box to initiate phylogenetic inference using PhyML (Fig. 9). The location of PhyML.exe will need to be added at first use. Upon completion a dialog box will open with links that will open the resulting tree in Mesquite for visualization. Mesquite (https://www.mesquiteproject.org/; Maddison and Maddison 2026) can also be used to label foreground branches/clades for the PAML branch, branch-site, and clade models implemented in the Selection Analyses (PAML) module (see PAML manual for details). Results will be automatically saved in the selected working directory. A complete history of BLASTN, MUSCLE, PRANK, and PhyML jobs is stored and can be accessed from the Gene Sequences dropdown menu.

**Figure 8.**
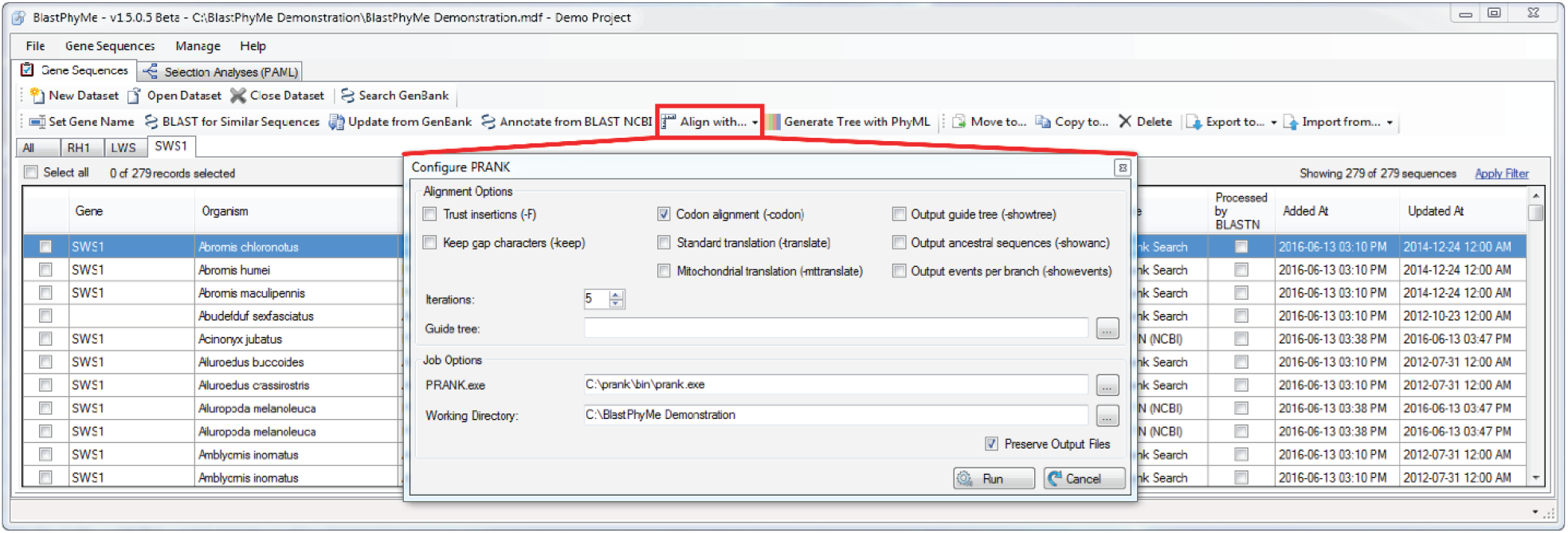
PRANK alignment window. A similar window is also available to set up alignments using MUSCLE.

**Figure 9.**
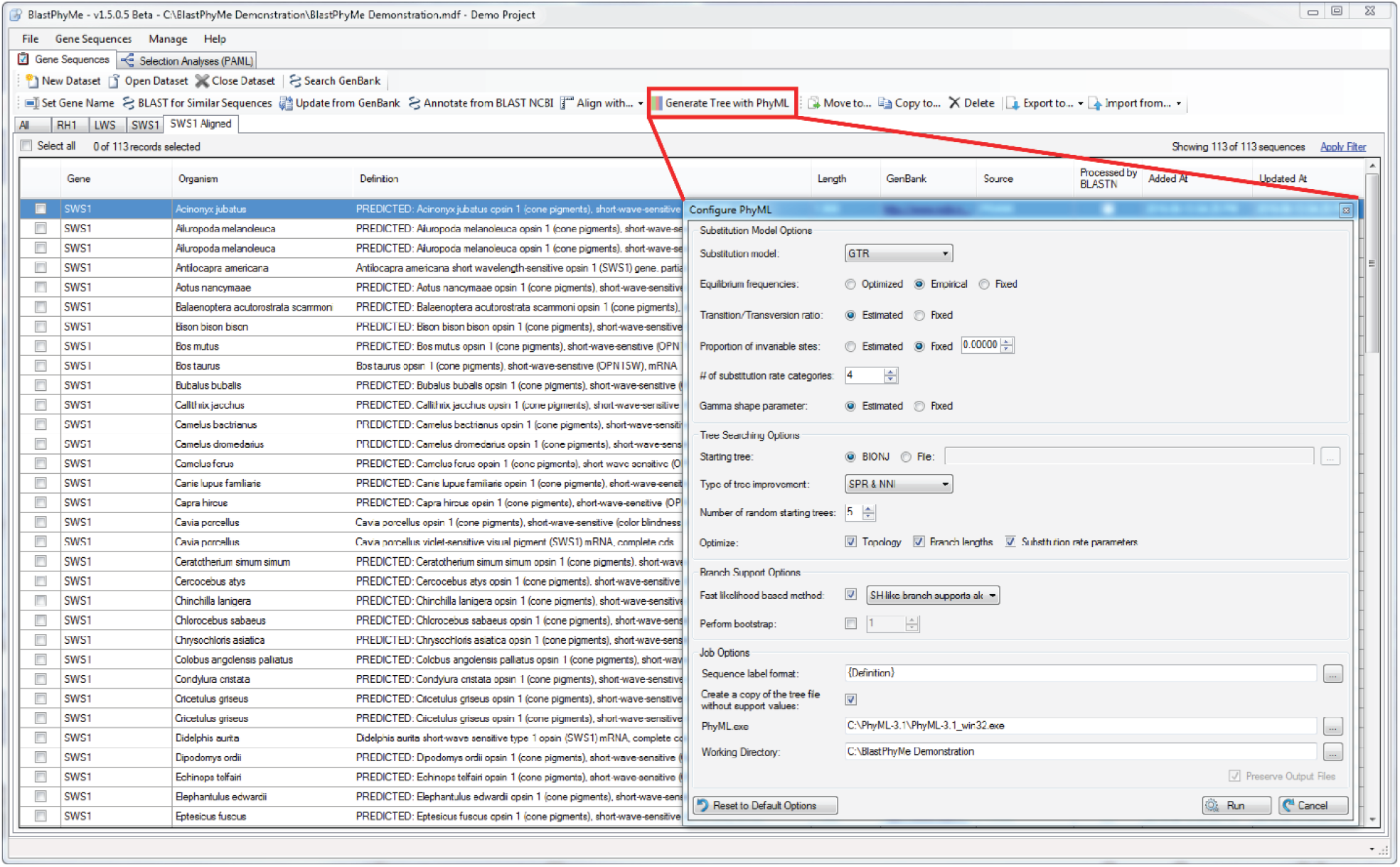
PhyML window to set up a phylogenetic analysis.

### Selection Analyses (PAML)

The Selection Analyses (PAML) module is similarly organized into datasets except that instead of storing sequences and GenBank records they store the results of PAML analyses. PAML jobs are initiated using the ‘New PAML Job’ button. This will open a dialog box allowing PAML analyses to be set-up (Fig. 10). Upon first use the location of codeml.exe will need to be specified. The number of processes to use and the working (output) directory should also be specified.

**Figure 10.**
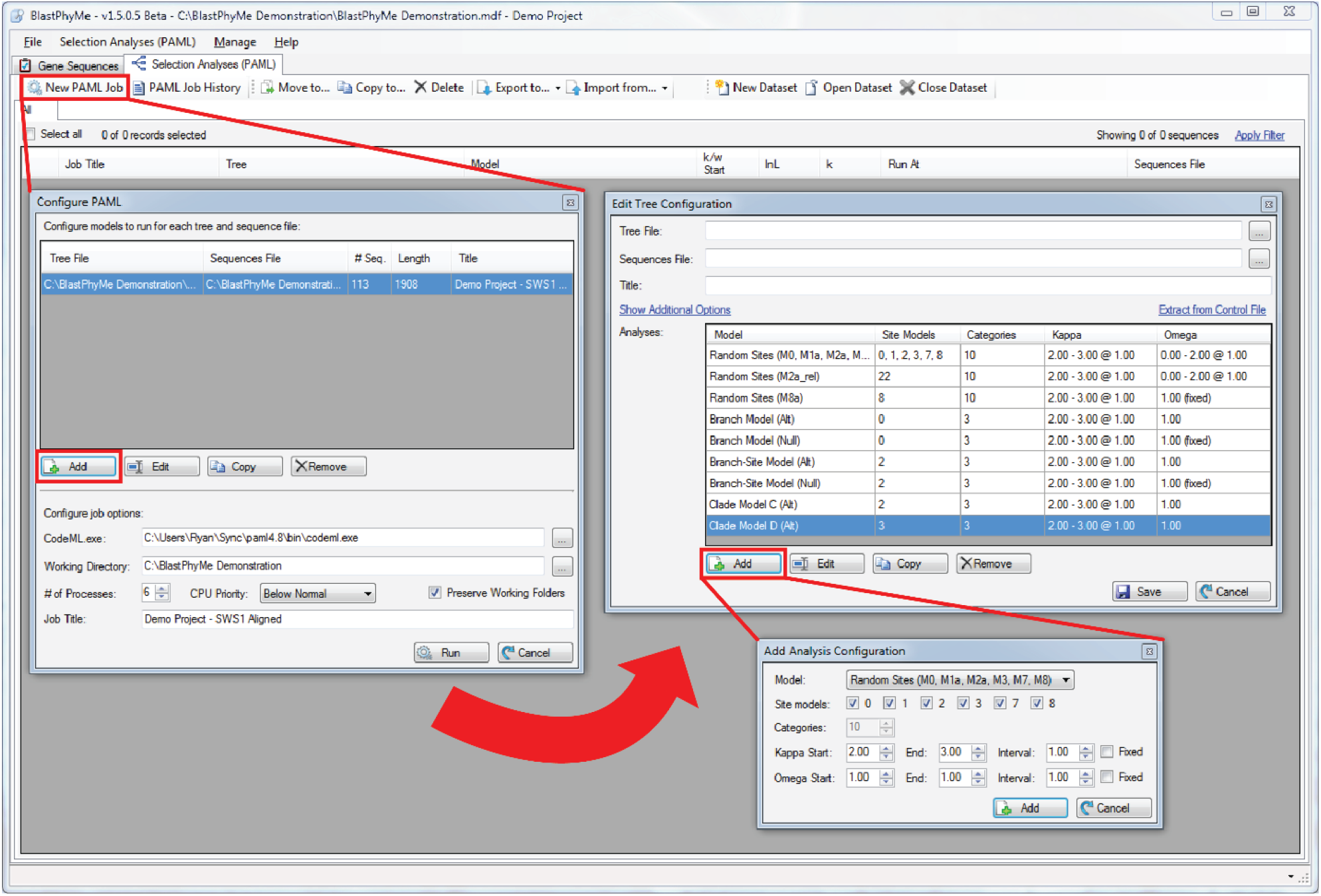
Interfaces used to set up and run PAML jobs.

**Figure 11.**
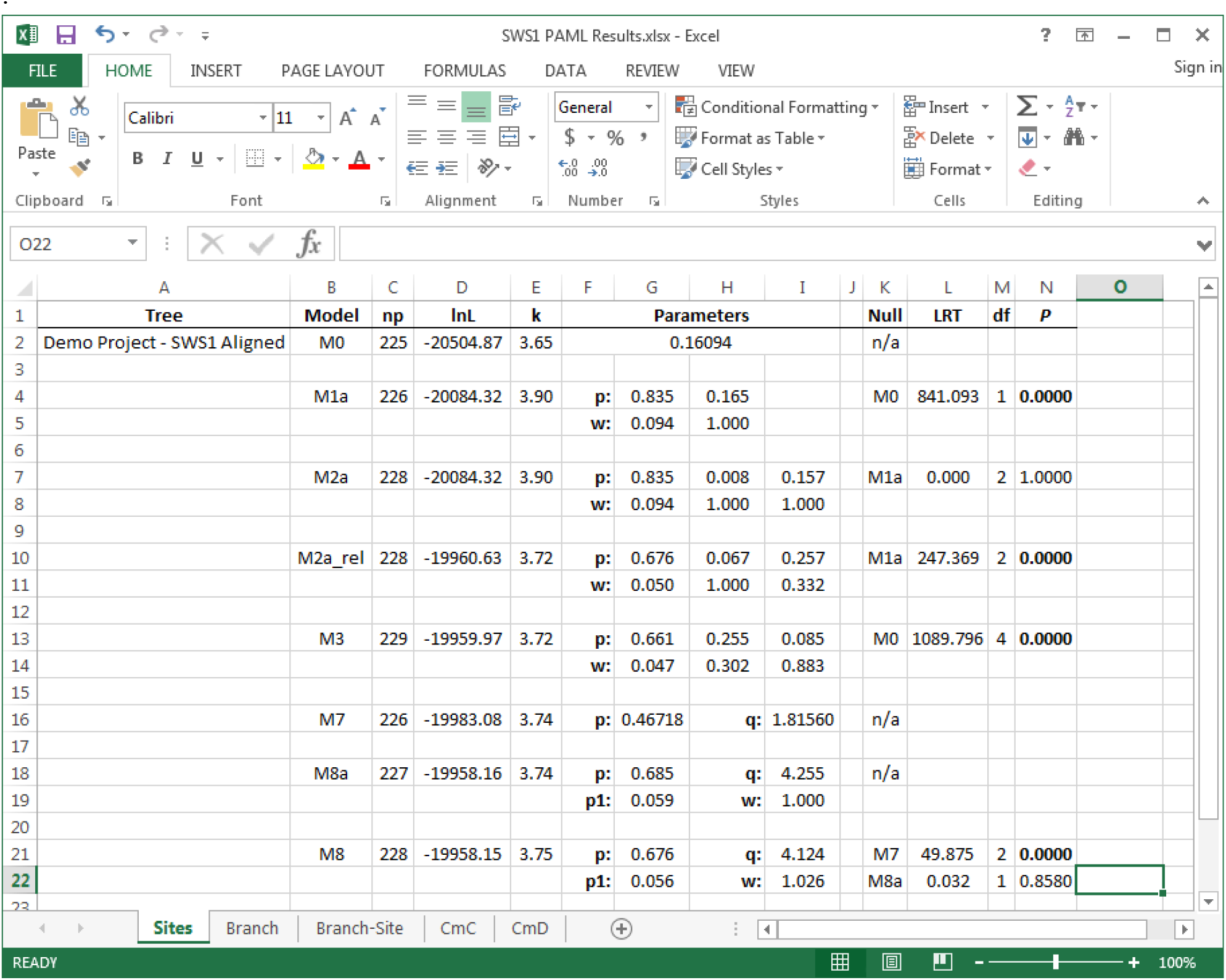
Excel table exported from BlastPhyMe using the “Export to” function. The table is automatically organized and formatted, with the appropriate likelihood-ratio test statistics and P values calculated between models.

When creating a new PAML job, clicking the ‘Add’ button will open a new dialog box (Fig. 10). Here a tree and sequence alignment file will need to be specified. If a phylogenetic analysis was performed using BlastPhyMe these can be found in the PhyML output folder that was specified. The ‘Add’ button in this dialog window allows models to be specified to run with the selected tree and alignment files. Models are specified using the drop-down menu, and for the site models multiple models can be specified simultaneously using the check boxes (Note: completion time is often faster when site models are added individually). Starting values for kappa and omega can be set and specifying a range of values will automatically set-up replicated analyses with each combination of starting values in the range. Multiple sets of models can be run for each tree and alignment pair and multiple tree and alignment pairs can be run with each PAML Job. Once executed with the ‘Run’ button, BlastPhyMe will initiate codeml for the specified number of processes and will run sequentially through each specified model until complete. A progress window will be displayed, but this can be closed and the job will continue to be run in the background.

Upon completion of a PAML job a results dialog box will appear. Each line will display the best replicate for the specified model. These can be double clicked to see the results of all replicates. Selected results can be added to a new or existing dataset. From within a dataset selected results can be exported to a preformatted Excel table using the ‘Export to’ function (Fig. 10). This produces a publication-ready table that automatically computes the appropriate likelihood ratio tests to determine significance.

## Example of Use

BlastPhyMe has been applied in published, peer-reviewed molecular evolution research, demonstrating the software’s ability for large-scale comparative selection analyses. Boyette et al. (2024) used the software to run codeml-based selection analysis on nonvisual opsin genes across 102 frog species spanning 34 families, applying PAML’s random-site models to estimate rates of nonsynonymous to synonymous nucleotide substitutions (ω or dN/dS) for inferring alignment-wide selection patterns and testing for positive selection acting on nonvisual opsin genes. The clade model was also used to test divergent selective pressures across different lineages.

## Competitive Landscape

Several existing tools leverage the codeml framework for selection analysis, including IDEA (Simons et al., 2008), IMPACT_S (Maldonado et al., 2014), LMAP (Maldonado et al., 2016), eto-evol (Huerta-Cepas et al., 2016), SlimCodeML (Schabauer et al., 2012), and VESPA (Webb et al., 2017), each differing in interface type, supported codon models, automated LRT capabilities, and multithreading support. Three tools were highlighted for direct comparison. BASE (Forni et al., 2021) uses a command-line (BASH and R) interface, employs Transeq for stop codon checking, performs branch length optimization and gene-tree inference via RAxML, and uses the Phangorn R package for tree annotation; its key distinguishing feature is the ability to integrate nonubiquitous genes by restricting analysis to genes present in the branch/clade of interest or applying a threshold for missing species. paPAML (Steffen et al., 2022) offers parallel processing of codeml and HyPhy analyses across multiple cores, a custom CDS-extraction tool built with Biopython, support for detecting selection in prokaryotic genes, and negative selection testing, though it is recommended for UNIX-based systems with multiple processors. EasyCodeML (Gao et al., 2019) provides a cross-platform Java-based GUI with one-click branch labeling, drag-and-drop file import, both preset and custom analysis modes, and a built-in sequence format converter.

BlastPhyMe distinguishes itself from these competing tools through several combined features: comprehensive support for the full range of codon models (branch, branch-site, and clade models), which are often limited or unavailable in competing software; a portable database framework (.mdf file) for efficient sequence data organization and management; integration of additional bioinformatics tools such as NCBI BLAST search and PhyML within a single application, eliminating the need for separate processing steps required by similar tools; storage of complete job histories to support reproducibility; automated parsing of codeml output into pre-formatted Excel spreadsheets with likelihood ratio tests computed automatically; and an intuitive GUI that removes the need for scripting, broadening accessibility of phylogenetic selection analysis to a wider range of researchers.

## Conclusions

BlastPhyMe saves researchers of all bioinformatics experience levels considerable time when generating and analyzing sequence datasets. BlastPhyMe incorporates existing services and utilities in a novel portable database framework with intuitive user interfaces not offered by other bioinformatics programs. BlastPhyMe currently runs on Windows 10 and 11 and an active internet connection is required only for functions that communicate with NCBI web services. BlastPhyMe is still under active development with planned updates to expand the available phylogenetic and molecular-evolutionary analyses.

## Acknowledgements

Development of this software was supported by NSERC Discovery Grants (to RKS and BSWC).

